# Vat Photopolymerization of Porous Scaffolds: Stabilization and Layer Thickness Control for Micron-Scale Accuracy

**DOI:** 10.1101/2025.01.07.631751

**Authors:** Guoyao Chen, Buddy D. Ratner

## Abstract

Vat Photopolymerization (VPP) holds much promise for producing biomaterial constructs such as porous scaffolds. However, achieving micron-scale pore dimensions with precision presents a challenge. This study offers an innovative approach to stabilize the silicone elastomer vat surface permitting micron-scale layer thickness accuracy to be maintained. Internal and surface contamination on the poly(dimethyl siloxane) (PDMS) vat surface were observed and effectively controlled with a pre-saturation methodology, and porous structures with cubical pores were then printed with varying layer thicknesses. These structures demonstrate the ability to achieve micrometric resolution and layer thicknesses as fine as 32 µm. A scaffold suitable for *in vivo* implantation with 40 µm cubical pores was successfully printed within 5 hours using a stabilized PDMS vat surface. Additionally, the methodology’s adaptability to intricate non-linear edge porous structures underscores its versatility across X, Y, and Z-axis.

## 1. Introduction

Three-dimensional (3D) printing, also known as additive manufacturing (AM), or rapid prototyping (RP), is a potentially powerful manufacturing method and is recognized as a versatile tool for precise fabrication of various devices in many fields including medicine[1-3], wastewater treatment[4, 5], energy storage[6, 7], microfluidics[8-10] and material science[11, 12]. 3D printing can produce complex architecture following a computer-generated design and can be fabricated using a diverse array of materials including polymers[13-15], metals[16-18], bio-inks[19], and ceramics[20]. This has led to the development of this technology for biomedical applications in both research and bio-industrial settings. The growing demand for customized pharmaceuticals and medical devices increases the impact of 3D printing, along with the demand for fast and high-resolution manufacturing processes[21].

To date, primary 3D printing techniques used in the biomedical field are vat photopolymerization (VPP) based stereolithography (SLA)[22-24], two-photon polymerization (TPP)[25], digital light processing (DLP)[26], continuous liquid interface production (CLIP)[27] and extrusion-based systems (polyjet, ink-jet and fused deposition modeling, FDM)[28, 29]. Resolutions range from 1 µm to 500 µm with some methods providing high spatial resolution (<50 µm)[30, 31]. The majority of these techniques are unable to achieve micron-level resolution when printing porous scaffolds, primarily due to challenges in accurately controlling pore dimensions along the Z-axis.

Porous scaffolds play a critical role in tissue engineering and regenerative medicine applications by providing a supportive structure for tissue regeneration[32]. The shape and size of the pores in the scaffold are crucial in determining its biocompatibility, mechanical properties, and cell behavior[33]. Spherical pores have been used in porous scaffolds, such as the precision porous scaffold with all pores approximately 40 µm in diameter invented at the University of Washington that show the ability to reduce or eliminate the fibrous capsule associated with the foreign body reaction[34, 35]. However, for interconnected, spherical pores of <100 µm, fabricating the scaffold using 3D printing becomes challenging, primarily due to over-curing issues. A possible approach for utilizing 3D printing to manufacture such high spatial resolution porous scaffolds is to use cubical pores.

Cubical pore shapes offer an alternative structure for tissue engineering scaffolds and can be more easily fabricated with 3D printing than scaffolds with spherical, interconnected pores. Cubical pores offer anisotropic mechanical properties that are desirable for specific applications, such as bone tissue engineering[36]. In 2018, Accardo et al.[37] reported the KLOE 3D lithographic laser system utilizing the SLA technique, capable of producing meso-scale structures. This system has the potential for printing scaffolds featuring micron-scale interconnected pores, for example 40 µm cubical pores.

SLA, as one of the earliest and most well-known forms of VPP techniques, is regarded as a rapid prototyping process and was developed in the late 1980s[38]. Anti-stick films or coatings are employed on the vat surfaces of certain VPP printers, including some SLA printers, to ensure that each newly fabricated layer adheres to the previously printed part rather than the vat surface, thereby facilitating a smooth fabrication process[39, 40]. This ensures that the VPP printer maintains relatively high speed while preserving high-resolution printing. Poly(dimethyl siloxane) (PDMS), Polytetrafluoroethylene (PTFE), and Fluorinated Ethylene Propylene (FEP) are commonly used as anti-stick materials, while PDMS is commonly used as a coating material due to its oxygen absorption properties, PTFE and FEP are frequently employed as air-permeable films that facilitate oxygen permeation[41, 42]. The oxygen present on their surfaces acts as a photo-inhibitor during the fabrication process, preventing the fabricated structure from adhering to the vat surface[43, 44]. The KLOE 3D lithographic laser system mentioned above utilizes PDMS as the vat coating, same as multiple SLA printers. SLA uses a .stl file to interpret a computer-aided design file that is communicated electronically to the 3D printer. Photoinitiation triggers monomers and oligomers to cross-link to form polymer structures[45]. Compared with other 3D printing methods, while most fabrication techniques have a resolution of 50–200 µm, many commercially available stereolithography 3D printers can build objects at an accuracy of 20 µm or lower[46]. In the Z-direction, the layer thickness can reach a resolution at 50 µm[47], but over-curing is an obstacle in maintaining the layer thickness, a challenge for fabricating high-resolution, multi-layer structures.

With an emphasis on addressing the over-curing issue in the Z-direction to ensure accurate cavity printing and establishing a stable printing environment with control over layer thickness for the production of scaffolds featuring cubic pores, this study presents a method for stabilizing SLA 3D printing performance. The KLOE high-resolution 3D lithographic laser system is used as a case study. We designed and integrated two distinct cubic porous units into diverse structures, aiming to create implantable-sized, high spatial resolution porous scaffolds amenable to 3D printing, enabling future *in vivo* investigations.

## 2. Material and methods

### 2.1 Material

A commercial acrylic-based 3D printing resin, VITRA DS2000 (DWS, lot# 10200075), was utilized to manufacture all the designs and experiments presented in this study. To create the PDMS print surfaces used in the monomer vat, a Sylgard-184 silicone elastomer kit (Dow Chemical Co.) was employed.

### 2.2 Sensitivity test

To assess the performance achieved with different pairings of laser intensity and laser writing velocity, a test pattern sensitivity test developed by KLOE was used [1.5 mm(X) × 1.5 mm(Y) base, 63 (7×9) 100 µm × 100 µm (X, Y)). The test pattern consists of 63 squares assessing 7 different print head velocities and 9 different laser intensities – the printer modulations. A 100 µm × 100 µm (X, Y) square at the top left corner was used as a direction reference. All the reported sensitivity test results are performed with a layer thickness 64 µm, 7 velocities (100 mm/s, 80 mm/s, 60 mm/s, 50 mm/s, 40 mm/s, 30 mm/s, 20 mm/s), the 9 modulations (100%, 80%, 60%, 50%, 40%, 30%, 20%, 10%, 5%). The reference squares were printed with 15 mm/s velocity and 50% modulation. After the tests were printed, they were immersed in two subsequent isopropanol rinses for 10 minutes to remove additional resin attached to the surface and dried at room temperature for 10 minutes.

### 2.3 Design of porous scaffolds

Porous scaffolds were fabricated using two distinct unit cells: the cubical architecture unit and the “cube-in-cube” architecture unit, here also refer as the MS unit resembling the Menger Sponge structure. The cylinder-shaped porous scaffolds measure 5 mm (X) × 5 mm (Y) × 2 mm (Z) in overall size. Additionally, the UW-shaped structure has dimensions of 8 mm (X) × 4 mm (Y) × 0.5 mm (Z), while the CDI-shaped structure measures 8 mm (X) × 3 mm (Y) × 0.5 mm (Z). Detailed schematic diagrams are provided in the discussion section.

### 2.4 3D printing of porous scaffolds

The sensitivity test pattern and the 3D scaffolds were printed using the KLOE high resolution 3D lithographic laser system, an SLA layer-by-layer 3D printer with a 375 nm laser. After the samples were printed, they were immersed in 2 isopropanol rinses for 10 minutes and 24 hours to remove uncured resin attached to the surface, and then dried at room temperature for 24 hours. For complex structures, due to the tortuous porosity, a longer post washing time is needed.

### 2.5 Scanning electron microscopy (SEM)

SEM images were acquired on the printed samples and test specimens after sputtering with gold using a SNE-3200M Scanning Electron Microscope (SEM) with an acceleration voltage at 5 kV.

### 2.6 PDMS coating on the vat printing surface

A 12g mixture of the Sylgard-184 silicone elastomer, consisting of a 10:1 ratio of base polymer to cross-linker, was used to coat the polymerization vat and form a layer of PDMS which absorbs/transmits oxygen and inhibits polymerization at the interface during structure fabrication. This ensures that the polymer in contact with the PDMS does not polymerize during the printing process, allowing the printing stage to move up with the polymerized part.

### 2.7 Testing for PDMS stabilization

A total of 3 g of Sylgard-184 silicone elastomer, with a mixing ratio of 10:1 between the base polymer and cross-linker, was employed to create a layer of PDMS on individual wells of a 6-well plate. This procedure was repeated 20 times, with every 5 wells forming a set, resulting in a total of 4 sets (n = 4). Following the PDMS coating for each set of wells, 5 g of DS2000 was applied to 4 out of 5 wells in each set, and left for durations of 10 minutes, 60 minutes, 24 hours, and 120 hours, respectively; the remaining well served as the blank control. Afterward, the DS2000 was removed, and the wells were washed with isopropanol until residual viscous liquid was no longer visible. Subsequently, the wells were placed in a Spectrolinker XL-1500 UV Crosslinker, which was equipped with 365 nm UV lights (BLE-1T151), for a duration of 9 minutes. Finally, the outcomes of each set were compared using the BioTek multi-mode plate reader with a 400 nm blue filter.

### 2.8 PDMS stabilization process

The PDMS coating was immersed in 40 mL DS2000 for 120 hours after a sensitivity test was performed. The liquid resin was then removed, and the PDMS was soaked in isopropanol for 30 minutes, followed by a 5-minute rinse with isopropanol until no liquid resin can be seen. The vat was then placed inside a UV chamber (Spectrolinker XL-1500 UV Crosslinker) with 365 nm UV lights (BLE-1T151) for 9 minutes. The above steps were repeated until the sensitivity test results stabilized within a range of 42-49 printed squares. A detailed discussion of this aspect and the rationale for this range will be provided in Section 3.1.

### 2.9 Measuring the layer thickness and pore size

The software ImageJ is employed for assessing the layer thickness and pore sizes of the scaffolds. In cases where edges appear straight and distinct, the original scale bar featured in the SEM image is utilized as the reference. However, certain scaffolds exhibit non-linear edges, leading to a perspective effect in the 2D SEM images. Consequently, accurately measuring certain pore sizes using existing SEM scales becomes challenging. To overcome this issue, a new measurement technique is introduced. In proximity to the pores requiring measurement, two parallel lines are randomly selected in the Z-direction, typically along the edges of each layer. Given the known layer thickness, the distance between these reference lines is employed to establish a new scale for pore size measurement. All subsequent pore size data are determined using this approach. To improve data precision, only Z-direction distances are taken into account, and for pores within the same layer, four random measurements are conducted (n = 4), as shown in Figure 1 (a).

**Figure 1.**
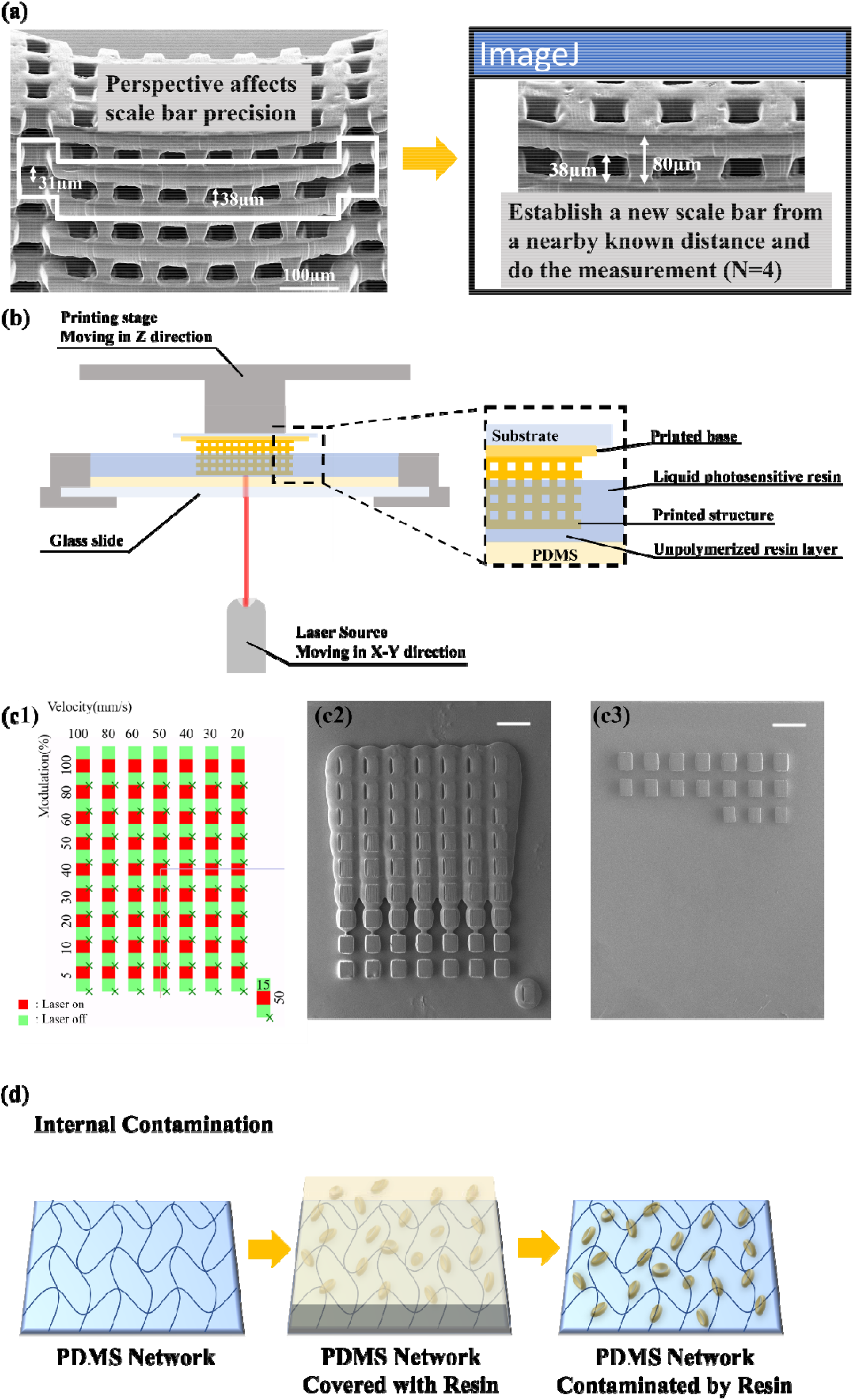
(a) schematic for perspective-adjusted measurements with ImageJ, (b) schematic diagram of the print stage of the KLOE SLA 3D printer, (c1) Schematic diagram of the sensitivity test, (c2) the result with a new PDMS surface (scale bar = 200 µm) and (c3) the result after repeated reuse of the PDMS (scale bar = 200 µm), and (d) schematic diagram suggesting internal contamination after repeated reuse of the PDMS layer

## 3. Results and Discussion

### 3.1 The PDMS print surface and sensitivity tests

The KLOE high resolution 3D lithographic laser system is utilized for fabricating all structures examined in this study. A diagrammatic representation of the printer is depicted in Figure 1 (b). Structures are generated in an inverted manner, with the printing stage moving along the Z-axis and the laser source operating within the X-Y plane. To ensure uniform photo polymerization, a base is initially printed before the main structure, unless otherwise stated. A layer of unpolymerized resin is always present between the printed structure and the PDMS due to oxygen absorbed by the PDMS, which hampers polymerization. It is evident that the PDMS surface significantly impacts the final printed layer, necessitating precise control of the PDMS surface to achieve accurate control of the actual layer thickness.

Prior to the fabrication of a structure, sensitivity tests were conducted to determine the optimal laser velocity and modulation (on and off time), illustrated in Figure 1(c1). It was observed that the test results would exhibit changes over time (as depicted in Figure 1(c2) and (c3)) when the PDMS surface is used repeatedly. This phenomenon arises regardless of the chosen printing parameters and is attributed to the resin penetrating the PDMS network and undergoing photocuring during the UV laser-based printing process (essentially a swelling effect), as depicted in Figure 1(d). This phenomenon is subsequently referred to as internal contamination.

A pre-saturation procedure might effectively address this issue. Saturating the PDMS network with DS2000 and subsequently exposing it to UV light before engaging in further procedures would ensure the stability of the PDMS print surface performance. This specific procedure is henceforth termed as the stabilization process, as illustrated in Figure 2(a). In order to optimize the conditions for achieving the ideal PDMS stabilization environment, experiments were required to determine the operational parameters of this process.

**Figure 2.**
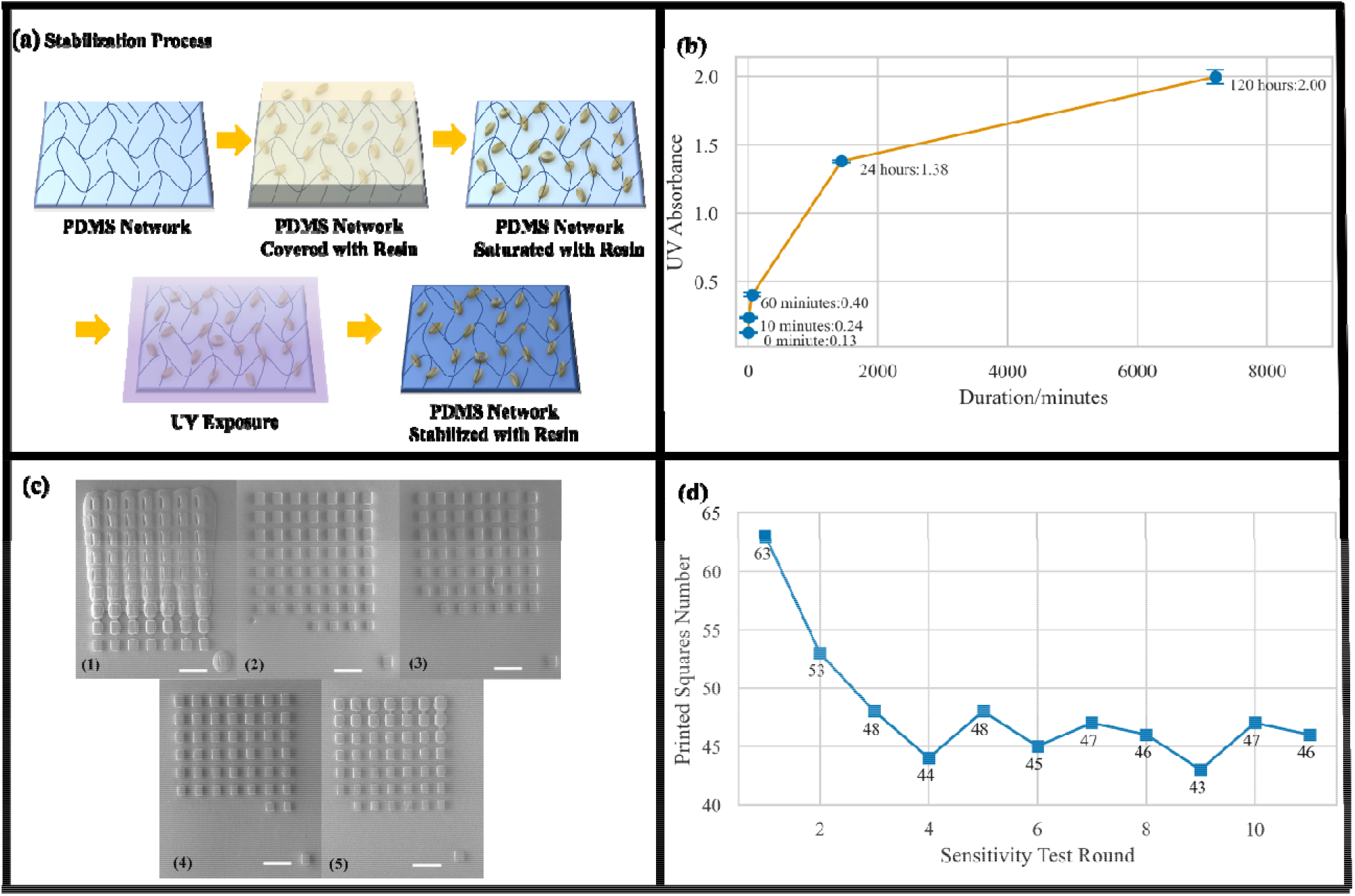
(a) schematic diagram of stabilization process of PDMS, (b) 365 nm UV absorbance result of PDMS coatings at varied DS2000 immersion times, (c) the first 5 sensitivity test results for the stabilization process (scalebar: 300 µm), and (d) the pattern of printed squares observed at 120-hour intervals stabilization (#1-#5) and randomly selected intervals stabilization (#6-#11), serves as evidence of the PDMS stability.

To address this concern, an initial investigation was undertaken involving small, trial-size PDMS sections that would fit into our plate reader UV spectrometer. The findings revealed increases in UV absorbance corresponding to length of immersion time, as illustrated in Figure 2(b) below. The rate of penetration declined over time, yet a small degree of penetration persisted between the 24-hour and 120-hour intervals. Consequently, the duration of 120 hours immersion in DS2000 was selected as the optimal penetration time for the stabilization process and was subsequently applied to the PDMS coating on the vat.

In contrast to the UV absorbance analysis used for smaller PDMS coating sections using a plate reader, the PDMS coating on the vat presents a challenge due to its larger size, making it unsuitable for plate reader measurements. To address this, the sensitivity test described earlier was employed to evaluate the outcomes. The investigation revealed that subjecting the PDMS coating to two rounds of 120-hour DS2000 immersions, followed by a 9-minute UV exposure to cure the absorbed DS2000, resulted in reasonably consistent and stable sensitivity test results. The threshold laser modulation discernible in the sensitivity test was approximately 20%. Thus, following the completion of the second round of the stabilization process, relatively stable outcomes were achieved, exhibiting a range of 42-49 printed squares. These initial five results are illustrated in Figure 2(c), and the trend of the outcomes is depicted in Figure 2(d), indicating a convergence following the stabilization process. We conclude that saturation of internal contamination on the PDMS print surface leads to predictable results. The variation in the number of printed squares may be attributed to the high printing velocity, inducing vibration that imparts a shear force on the printed squares, potentially causing less securely adhered squares to detach from the printing base.

This pre-saturation stabilization process should be widely applicable to any SLA printer utilizing a PDMS vat coating. By achieving a stable PDMS print surface, a majority of combinations of printing velocity and laser modulation exhibit favorable outcomes, enhancing our capability to attain precise layer thicknesses. Our prior findings indicate that the KLOE high-resolution 3D lithographic laser system can attain a resolution of 5 µm in the X-Y plane[48]. With the incorporation of a stabilized PDMS surface, the fabrication of porous scaffolds with micron-level resolution becomes feasible.

### 3.2 3D printed porous scaffolds with “cube-in-cube” units

The stabilized PDMS printing surface enables a precisely controlled layer thickness. This stabilized surface was used to fabricate porous scaffolds with cubical pores.

The “cube-in-cube” architectural unit, as illustrated in Figure 3(a) below, is composed of three distinct layers that can be segmented into printable slices. To clarify the architecture, a different view of the unit and a 2 × 2 × 2 assembly structure are also shown in Figure 3(a2) and (a3). This design emulates the precision-spherical pore structure described in the introductory section, albeit featuring cubical pore geometry instead of spherical. The objective is to align the fabrication process with the capabilities of the Kloe 3D printer. The interconnects, are 1/3 of the pore size analogous to spherical pore scaffold. The cubic architecture resembles the level 2 Menger Sponge (MS) structure [49, 50]. All the following scaffolds produced using the “cube-in-cube” unit will be denoted as MS scaffolds. In this study, MS scaffolds with interconnectivity of both 64 µm and 32 µm were manufactured.

**Figure 3.**
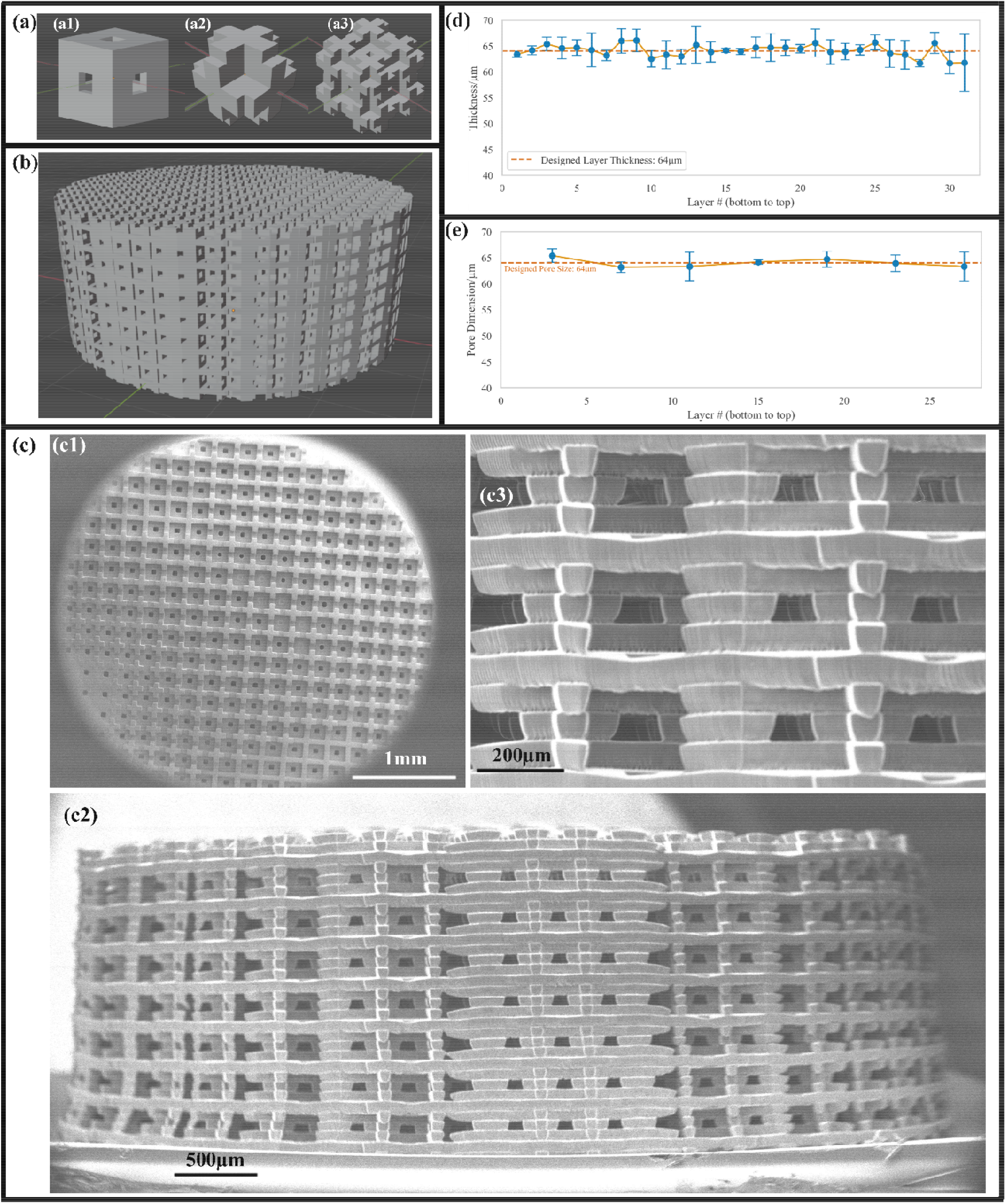
(a) the MS architecture unit, (a1) the original unit, (a2) the corner-view unit, and (a3) the 2 × 2 × 2 assembly structure of the corner-view unit, (b) a computer-generated image of 5mm(X) × 5mm(Y) × 2mm(Z) cylindrical-shaped MS scaffold with 64 µm interconnects, (c) SEM pictures of the printed MS scaffold with 64 µm interconnects (c1) the top view, (c2) the side view, and (c3) a zoom in view of the top layers showing the interconnectivity, (d) the layer thickness of the MS scaffold from bottom to top, and (e) the pore size of the MS scaffold from bottom to top

To assess the printer’s precision in producing layers with a thickness of 64 µm, we fabricated a cylindrical MS scaffold measuring 5mm (X) × 5mm (Y) × 2mm (Z) with 64 µm interconnects. The computer-generated representation can be found in Figure 3(b), while the SEM image of the actual, printed construct is displayed in Figure 3(c). The fabrication process began with a printing base created at 100% modulation and a velocity of 50 mm/s. Subsequently, the MS scaffold was manufactured using 40% modulation and a velocity of 60 mm/s, leading to a layer thickness of 64 µm. The entire fabrication process took approximately 3 hours. Given the scaffold’s high porosity, a substantial amount of uncured resin was trapped within the structure, necessitating post-printing rinsing. Consequently, the scaffold was subjected to two successive isopropanol rinses lasting 1 hour and 24 hours, respectively.

In Figure 3(c1), some of the interconnects are obstructed by polymerized resin. This could be attributed to resin retained within the scaffold, even after the isopropanol rinsing process and subsequently polymerized due to natural light exposure in the external environment.

The side view in Figure 3(c2) reveals the structural integrity maintained during the printing process. In Figure 3(c3), a closer view of the uppermost layers highlights the inner cubical pores of the scaffold, showcasing that most of the structure is uniform. Further insights into layer thickness and pore size measurements are presented in Figure 3d and 3e. It is evident that both the layer thickness and pore sizes across all layers closely align with the intended dimensions. This example effectively demonstrates that employing a stabilized vat coating and precise layer thickness control facilitates the rapid 3D printing of structures with repeatable layers and micron-level resolution. The scaffold exhibits a resolution between layers that manifests micrometric periodic features down to 64 µm. Importantly, the fabrication process is swift in comparison to two-photon polymerization additive manufacture[51].

To assess the printer’s precision in fabricating layers with a thickness of 32 µm, we fabricated a cylindrical-shaped MS scaffold with 32 µm interconnects, as depicted in Figure 4(a) (computer generated graphic) and showcased in Figure 4(b) (SEM results). The fabrication process commences with a printing base established at 100% modulation and a velocity of 50 mm/s. The construction of the cubical-porous scaffold ensues at 30% modulation and 60 mm/s, utilizing a 32 µm layer thickness, culminating in an overall fabrication duration of 6.5 hours.

**Figure 4.**
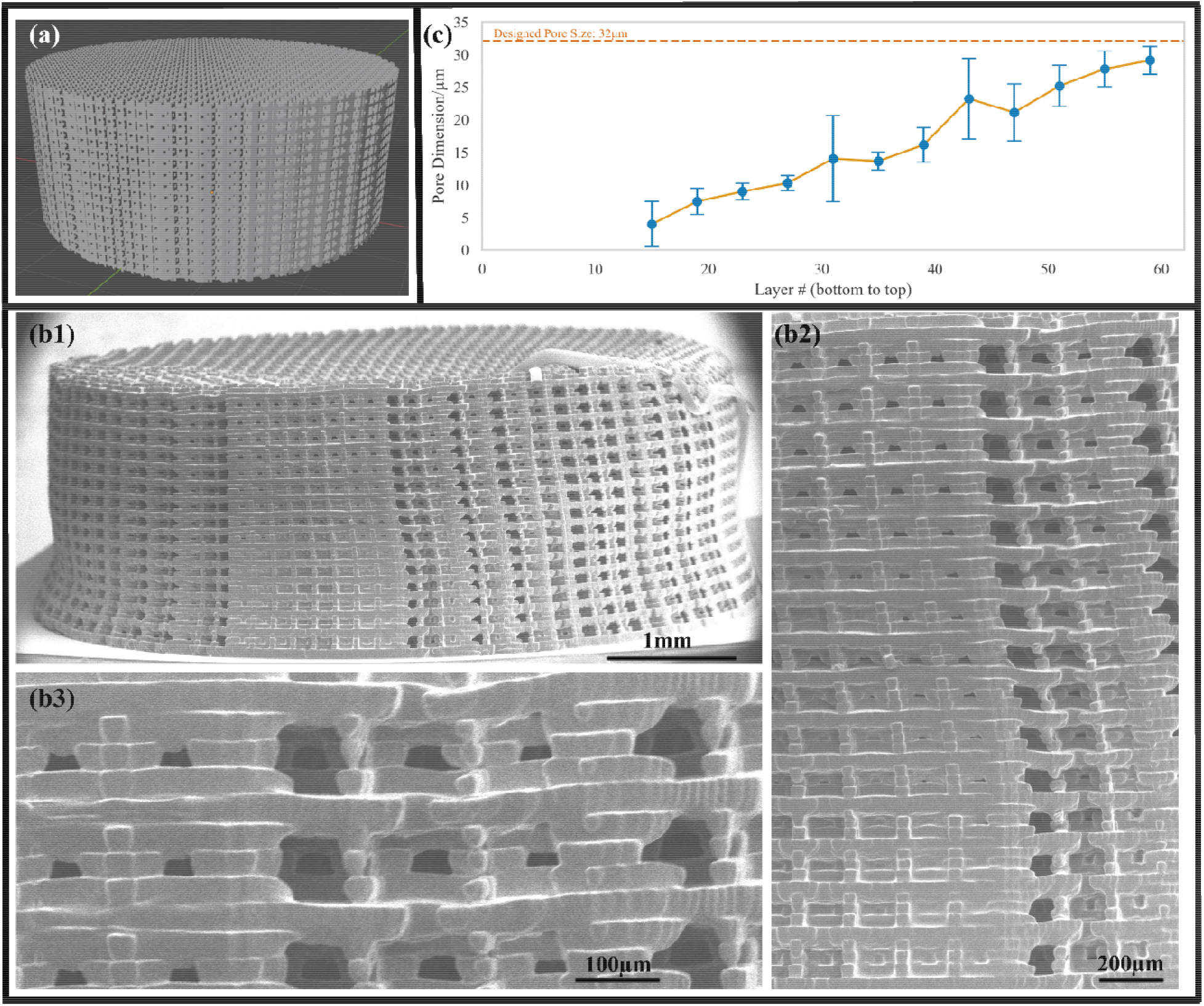
(a) computer-generated image of 5mm(X) × 5mm(Y) × 2mm(Z) cylindrical-shaped MS scaffold with 32 µm interconnects, (b) SEM pictures of the MS scaffold with 32 µm interconnects (b1) the side view, (b2) a zoomed-in view of the actual layer thickness change, and (b3) a zoomed-in view of the top layers showing the interconnectivity, and (c) the measured pore size of the MS scaffold with 32 µm interconnects from bottom to top

While the porosity geometry of this structure mirrors that illustrated in Figure 3(c), the interconnects are 12.5% of the prior design, necessitating an extended post-rinsing period. The sample underwent immersion in two isopropanol rinses, spanning 1 hour and 48 hours. Examination of the upper layers of the structure, detailed in Figure 4(b3), reveals an actual layer thickness down to 32 µm. However, Fig 4(b2) depicts variations in printed layer thickness along the printed height, with the initial layers surpassing 32 µm. This observation implies that the 30% modulation combined with a velocity of 60 mm/s fails to ensure consistent production of 32 µm thick layers. Furthermore, the analysis of measurable pore sizes in Figure 4(d) demonstrates fluctuations in the printing environment during the fabrication process, leading to alterations in the actual layer thickness over time.

After conducting tests with varying laser intensities spanning 15% to 40%, it became evident that maintaining a consistent layer thickness of 32 µm is challenging. Furthermore, employing a laser intensity below a certain threshold leads to incomplete shapes in the printed structure. Presently, a solution to this issue is not available, leaving the MS scaffold with 16 µm layers as a topic for future exploration. Notably, after the printing process, the area on the PDMS where the scaffold was created appears less transparent compared to the rest of the PDMS coatin, and the stain left by the printed structure can be wiped off with isopropanol after the printing process. This indicates an alteration in the surface condition during printing. This phenomenon is henceforth referred to as surface contamination.

Assuming the printer’s performance remains consistent, for each distinct structure, all layers would demand a uniform laser dose, i.e., the combination of laser intensity and velocity, to achieve an actual thickness approximating the intended value. If the laser dose surpasses the requisite level, over-curing occurs, resulting in the physical compression of the PDMS coating and the deposition of residual material, and the residual material would decrease the transparency of the PDMS. This decreased transparency effectively decreases actual laser dose leading to a thinner next layer. This phenomenon is evident in the cylindrical-shaped MS scaffold with 32 µm interconnects, as illustrated in Fig 4(b2).

Ideally, by optimizing the laser dose to achieve an appropriate value and targeting a layer thickness of around 32 µm, successful fabrication can be achieved. However, our findings suggest that excessively low laser doses hinder the integrity of the printed structure. A hypothesis emerges that during polymerization, insufficient laser energy prevents complete solidification of the resin, resulting in a semi-solid state with elevated viscosity. Such layers are prone to leaving residues on the PDMS coating during the formation process, leading to increased surface roughness and opacity. This enhanced roughness facilitates adhesion to PDMS during layer-by-layer polymerization, generating a cycle of residue accumulation that significantly impacts the overall printed structure.

Following this hypothesis, if the optimized laser dose for a particular layer thickness falls below the minimum required level for full solidification, the printed structure experiences variable layer thickness along the Z-axis and cannot be accurately produced, as demonstrated in Figure 4. In essence, there exists a theoretically achievable minimum layer thickness that can be tightly controlled, and 32 µm falls below this threshold. Guided by this hypothesis and considering our original goal of fabricating 40 µm pore-sized scaffolds, a new geometry has been designed and fabricated to evaluate whether 40 µm exceeds the required minimum layer thickness.

### 3.3 3D printed porous structures with 40 µm cubical units

With the objective in mind to create a 40 µm geometric structure that might show superior healing upon implantation, a cubical unit incorporating a 40 µm cubical void was designed and subsequently printed, as depicted in Fig 5(a) below. This unit can be divided into two repeatable printable slices. In contrast to the MS unit, it lacks smaller interconnects, with the pore size serving as the interconnect dimension. While not the original intended architecture, this design serves as an effective means to assess the PDMS print surface performance and identify an optimal layer thickness for precise printer control.

**Figure 5.**
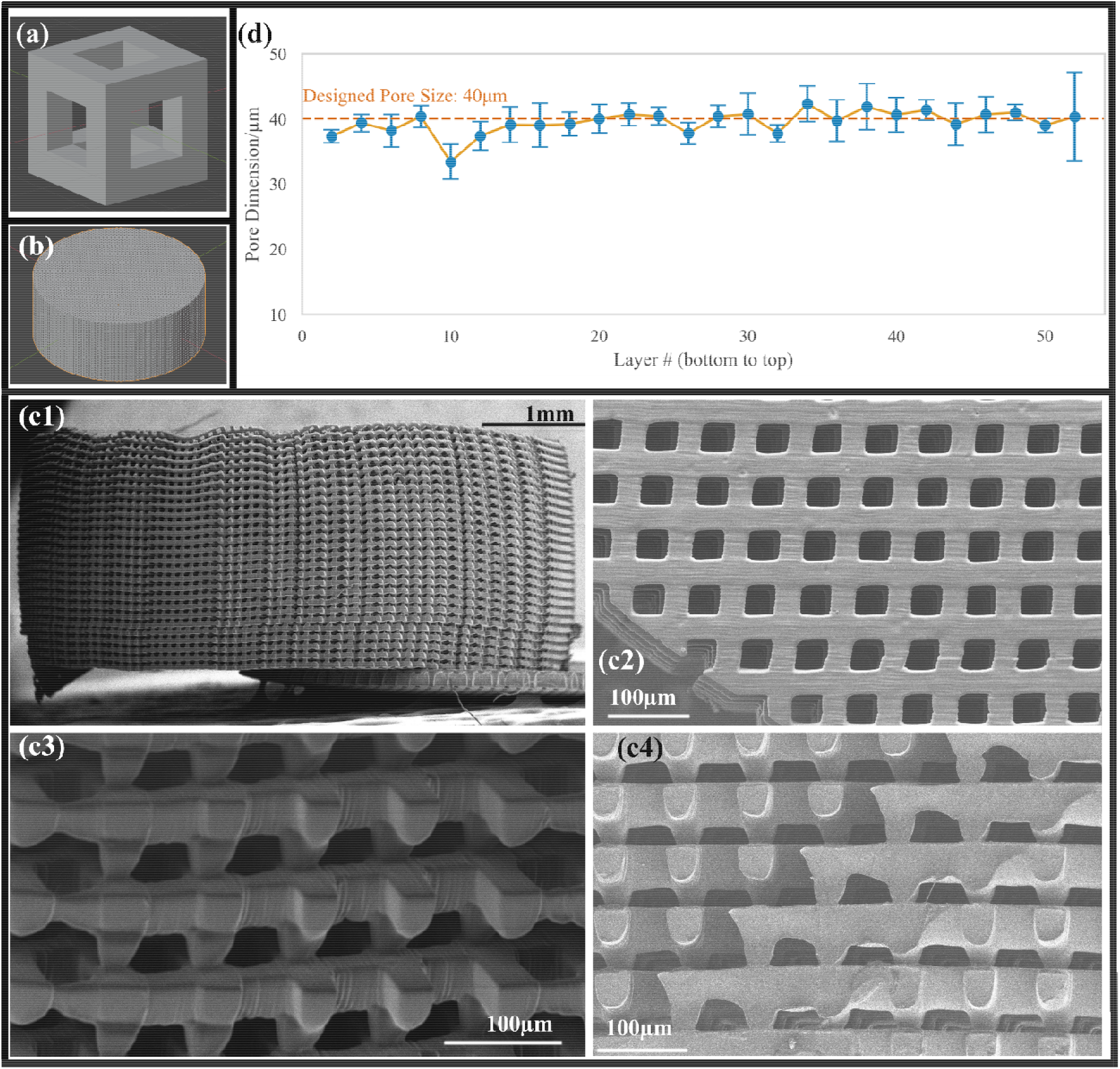
(a) computer-generated image of the cubical architecture unit, (b) computer-generated image of the cylinder-shaped cubical scaffold printed with 40 µm pore size, (c) SEM pictures of the square-disk shape cubical scaffold printed with 40 µm pore size, (c1) the side view, (c2) the corner of the top view, (c3) the zoom in side view of the top layers, (c4) the cross-section view after sectioning, and (d) the measured pore size of the cubical scaffold with 40 µm layer thickness from bottom to top

To assess the printer’s precision in fabricating 40 µm-thick layers, a cylindrical porous scaffold measuring 5mm (X) × 5mm (Y) × 2mm (Z) was created using cubical units, as depicted in Figures 5(a) and (b). The initial layer was used as the base and printed at a modulation of 50% and a velocity of 50 mm/s. The remaining scaffold was printed using a modulation of 35% and a velocity of 60 mm/s, resulting in a consistent layer thickness of 40 µm and a total fabrication time of 5 hours. Following printing, the scaffold underwent two consecutive isopropanol rinses lasting 1 hour and 24 hours. Illustrated in Figure 5(c1), the entire structure was successfully printed. Zoomed-in views, presented in Figures 5(c2) and (c3), provide detailed top and side perspectives of the scaffold, confirming accurate structure replication and pore dimensions as intended. As illustrated in Figure 5(c4), a cross-sectional view of the scaffold revealed that while the inner structures remained porous, slight over-curing in the middle caused the poresize to be smaller than the outer pores. In Figure 5(d), all pore sizes except for the 10th layer demonstrated consistent 40 µm dimensions, possibly attributed to volume shrinkage around that layer. Overall, the printed scaffold demonstrated the printer’s proficiency in generating 40-µm-thick layers, surpassing the minimum required layer thickness as hypothesized earlier.

The cubical porous configuration can be fabricated into larger-scale shapes that suggest the potential to fabricate real-world objects. As examples of the technological ability to create practical sized structures, Figures 6 (a1), (b1), and (c1) show SEM images of a UW-shaped structure, a CDI-shaped structure, and a vertically oriented CDI-shaped structure. The foundation (base) components were printed using a modulation of 100% and a velocity of 50 mm/s, while the remaining sections of the structures were printed using a modulation of 31% and a velocity of 60 mm/s. The fabrication durations were 69 minutes, 109 minutes, and 180 minutes and the corresponding outcomes are illustrated in Figure 6 (a), (b), and (c), respectively.

**Figure 6.**
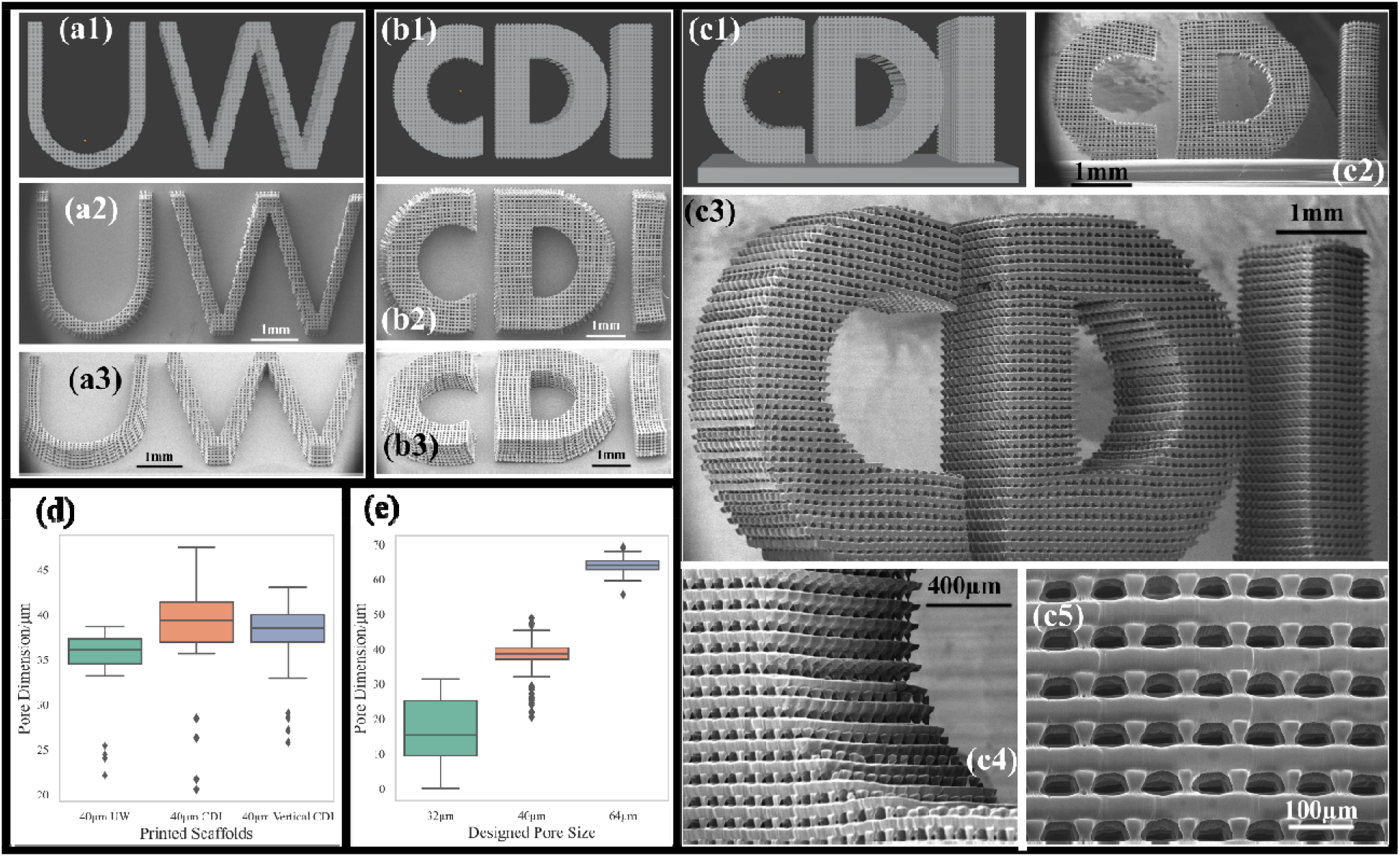
Printing of 3 complex structures with cubical pores, (a) UW-shaped porous scaffold(a1) computer-generated image of the UW-shaped structure with thin walls, (a2) top view of the result, (a3) tilted angle view, (b) horizontal CDI-shaped porous scaffold (b1) computer-generated image of the horizontal CDI-shaped structure with thick walls, (b2) top view of the result, (b3) tilted angle view, (c) vertical CDI-shaped porous scaffold (c1) computer-generated image of the vertical CDI-shaped structure, (c2) side view, (c3) tilted angle side view, (c4) high magnification view of the corner of the letter C, (c5) high magnification view of the letter I showing the pore size, (d) Box plot for cubical pore size distribution of these 3 structures, and (e) Box plot for cubical pore size distribution of different printing layer thicknesses

The CDI-shaped and UW-shaped structures were employed to assess the printer’s capability in maintaining layer thickness stability during the fabrication of intricate structures with complex X-Y plane edges. The UW-shaped structure features narrow walls, while the CDI-shaped structure incorporates thicker walls. It is noteworthy that, as depicted in both Fig. 6(b2) and (b3), the letter “I” within the CDI-shaped structure displays slight distortion due to pillars that were not properly attached between layers during the printing process. These unattached pillars were subsequently removed in the washing step, resulting in deformation. However, both the UW-shaped and CDI-shaped structures showcase the printer’s ability to produce intricate designs with non-linear X-Y plane edges using a layer thickness of 40 µm. The absence of substantial surface contamination under these printing conditions indicates that the laser dosage has been optimized to mitigate the impact of surface contaminants.

To demonstrate the printer’s ability to produce porous structures with non-linear Z-axis edges, the CDI-shaped structure was rotated by 90 degrees and printed vertically, as shown in Figure 6(c). This outcome affirms the printer’s competence in fabricating intricate porous scaffolds in all three dimensions. The outcomes presented in Figure 6(c4) reveal that the printer’s precision in printing pillar-like forms with controlled layer thickness can lead to the transformation of the initial rectangular cube design into trapezoidal or triangular side surfaces. This phenomenon is attributed to the Gaussian nature of the laser beam utilized in SLA printers. The narrow wave width of the laser beam, employed to prevent over-curing and control layer thickness, might result in slightly smaller actual sizes at the upper end of a given layer compared to the lower end. This effect contributes to the parabolic shape observed, which may deviate from the intended cubical pore shape. While this highlights extreme control over the laser beam, the presence of partially detached pillars may lead to structural deformations and collapse. Hence, a controlled degree of over-curing becomes essential to ensure overall structural stability.

From Figure 6(d), it is evident that while the layer thickness remains relatively consistent across the three structures, the actual pore size of the narrower walls, represented by the UW-shaped structure, is smaller compared to the thicker walls of the CDI-shaped structures. This discrepancy suggests that the UW-shaped structure, despite being printed with the same laser intensity, encounters over-curing issues. This phenomenon could be attributed to the fact that, with areas already polymerized by the laser, smaller structural dimensions will experience a higher frequency of laser exposure nearby. This increased frequency raises the likelihood of additional laser energy affecting those regions, leading to over-curing.

Consequently, even though the theoretical layer thickness may be the same across different structures, the required laser intensity varies due to their distinct dimensions. Through experimentation, we noted that the structures can be reprinted with a range of laser intensities and the result is stable. Remarkably, the actual layer thickness exhibits relatively small variation between layers, with no layer thickness fluctuations as shown in Figure 4(b3), suggesting that the surface contamination can be ignored in this instance.

The comparison of actual cubical pore size distribution, as depicted in Figure 6(e), reveals that under identical printing conditions, a 32 µm layer thickness does not consistently maintain the designated value. However, a modest 25% increase in the designed layer thickness to 40 µm significantly enhances stability. Despite some fluctuations in actual layer thickness under the same printing conditions, structural stability remains unaffected. Furthermore, elevating the layer thickness to 64 µm results in further improvement in stability.

Currently, our research endeavors are focused on investigating printer stability and reproducibility when printing structures with dimensions below 40 µm. Nevertheless, the viability of maintaining a layer thickness of 40 µm has been convincingly demonstrated through the successful printing of these three intricate structures.

## 4. Conclusion

The SLA 3D printer explored in this work (KLOE 3D lithographic laser system) has the potential to produce intricate multilayer structures with high precision and high resolution. However, methodological issues with the PDMS printing surface can degrade ultimate outcomes. We present a novel approach to achieve stability in the PDMS printing surface for SLA fabrication, effectively controlling two types of contamination during the printing process: internal and surface contamination. These proposed methods are compatible with most SLA printers and contribute to optimizing performance. Using our improved print surface pretreatment method, we successfully created two cylindrical-shaped porous scaffolds featuring Menger Sponge units, one with 32 µm interconnects and the other with 64 µm interconnects. Our results demonstrate the potential for achieving micrometric periodic features down to 64 µm and actual layer thicknesses approaching 32 µm using SLA techniques. It should be noted that there is a significant fabrication problem when layer thickness is reduced to 32 µm.

This study has effectively produced a cylinder-shaped scaffold with 40 µm cubical pores, showcasing the precise fabrication of slices with 40 µm thickness through the application of the stabilized PDMS coating. Although distinct from the precision-spherical pore structure, this scaffold design presents potential applications in material science, biomedical engineering, and device design. The stabilized PDMS coating enables the fabrication of intricate architectures with non-linear edges across all three spatial dimensions (X, Y, and Z) using a 40 µm layer thickness. Both the UW-shaped and CDI-shaped models were successfully printed, exhibiting structural robustness during printing.

In conclusion, our methodology allows for the precise creation of porous scaffolds with micron-level resolution, offering sizes and shapes suitable for animal implantation. This achievement opens potential applications of these scaffolds in fields such as material science, biomedical engineering, and device design. Furthermore, an *in vivo* study in mice has been conducted to evaluate the cellular response to the scaffold with 40 µm cubical pores. We look forward to sharing these results in our next paper, where we will compare cellular performance across different pore geometries.

## CRediT authorship contribution statement

**Guoyao Chen:** Investigation, Writing – original draft, Conceptualization, Methodology, Validation, Data Curation, Software, Writing – review & editing, Visualization, Supervision, Project administration. **Buddy Ratner:** Conceptualization, Writing – review & editing, Supervision, Funding acquisition.

## Acknowledgments

This work is funded by the University of Washington Center for Dialysis Innovation (CDI) which has been supported by the Northwest Kidney Centers.

